# Synthesis-aware generative design in trillion-scale chemical spaces for automated drug discovery

**DOI:** 10.64898/2026.10.01.755933

**Authors:** Seonghwan Seo, Yulseung Sung, Sang-Yeon Hwang, Mincheol Kang, Joonseong Lee, Samuele Bordi, Luka Raguž, Jihye Choi, Daniil Melnichenko, Wan Namkung, Sehan Lee, Jaechang Lim, Benedikt M. Wanner, Jung Min Han, Woo Youn Kim

**Affiliations:** Department of Chemistry, KAIST, Daejeon, Republic of Korea; Yonsei Institute of Pharmaceutical Sciences, College of Pharmacy, Yonsei University, Incheon, Republic of Korea; HITS Inc., Seoul, Republic of Korea; eMolecules Inc., Kemptthal, Switzerland; InnoCORE AI-CRED Institute, KAIST, Daejeon, Republic of Korea; Kim Jaechul Graduate School of Artificial Intelligence, KAIST, Daejeon, Republic of Korea

**Author notes:** Corresponding authors. (J.M.H.); (W.Y.K.). Division of Infectious Disease, Boston Children’s Hospital and Harvard Medical School, Boston, MA, USA. These authors contributed equally to this work (Seonghwan Seo, Yulseung Sung, Sang-Yeon Hwang).

## Abstract

Autonomous drug discovery via generative molecular design is critically bottlenecked by the production of chemically intractable structures. Multi-trillion-scale make-on-demand libraries guarantee synthetic feasibility, but conventional virtual screening cannot efficiently navigate these vast spaces or address multi-parameter optimization. We introduce Hyper Screening X, a generative AI framework integrating structure-based design with automated synthesis hardware. By encoding deterministic reaction logic in a generative flow network, the framework evaluates only 10 million candidates to navigate an 11-trillion-compound space, jointly optimizing physicochemical properties while ensuring compatibility with automated synthesis. Prospective validation against an SLC1A5 variant with a cryptic interface yielded a 96% synthesis success rate and a 50% functional hit rate, identifying two first-in-class lead compounds. This synthesis-aware strategy advances autonomous, closed-loop drug discovery.

## Introduction

Artificial intelligence (AI) is accelerating the shift toward autonomous scientific discovery (1–4). In drug discovery, this transition envisions a closed-loop design–make–test (DMT) cycle driven by generative AI for *de novo* molecular design (5–10). However, realizing a fully autonomous pipeline is hindered by three fundamental bottlenecks: data dependency, multi-parameter optimization (MPO), and synthetic feasibility. Because standard generative models need massive datasets of known active ligands (5, 11), the field has increasingly adopted structure-based drug design to bypass this data dependency using the 3D information of protein targets (12–14). The MPO process demands a strict balance between high potency and stringent physicochemical properties. Because these drug-like objectives frequently conflict, the probability of satisfying a comprehensive target product profile decays exponentially (15, 16), necessitating advanced AI to simultaneously co-optimize competing parameters across ultra-large chemical spaces. Although recent approaches incorporate synthesis-aware strategies using purchasable building blocks and reaction templates (17–25), these models still frequently produce synthetically intractable compounds or propose routes incompatible with automated synthesizers. Overcoming this challenge requires AI design guaranteed by synthesis directly coupled with automated hardware to unlock the vast chemical diversity essential for next-generation therapeutics.

To access this vast synthesizable chemical space, the field relies on the construction of ultra-large, make-on-demand combinatorial libraries navigated via virtual screening (VS) pipelines apart from generative approaches (26–28). While screening billion-scale spaces systematically improves the discovery of novel chemotypes (29–31), these billion-scale libraries frequently suffer from an optimization void, a null set lacking any molecule that meets the desired target product profile. Bridging this void requires access to trillion-scale chemical landscapes or beyond. However, extrapolating conventional VS to trillions of compounds is computationally prohibitive, leaving traditional methods unable to co-optimize multiple properties without extensive filtering.

Here, we introduce Hyper Screening X (HSX), a generative AI framework that overcomes these bottlenecks by shifting from static library screening to active, structure-based design within a fully machine-actionable space. We embed the deterministic reaction logic of make-on-demand combinatorial chemistry directly into a generative flow network (GFlowNet) (6, 32). Unlike standard reinforcement learning that frequently suffers from mode collapse, GFlowNets sample diverse, high-reward candidates proportionally to their objective distributions (7, 33). Recent pioneering works have shown the viability of coupling GFlowNets with synthesis pathways (20–22, 24, 34), with RxnFlow notably demonstrating their potential for large-scale chemical exploration (22). Building on these foundations, HSX represents a critical leap forward as the first framework to expand synthesis-aware generation and rigorous MPO across a multi-trillion-scale landscape. By formulating automated synthesis libraries (e.g., those by Synple Chem and eMolecules (35, 36)) as a structured decision tree (30), HSX efficiently navigates this vast space, simultaneously co-optimizing five or more conflicting physicochemical parameters while maintaining *>*80% synthetic feasibility.

To empirically demonstrate the capability of this synthesis-aware MPO in a highly challenging biological context, we validate this framework by targeting the mitochondrial glutamine transporter SLC1A5_var, a first-in-class target defined by a complex protein–protein interaction interface. Our *in silico* study shows that the HSX platform can accelerate MPO-driven drug design by approximately three orders of magnitude relative to standard VS workflows. Prospective experimental validation yielded a 50% hit rate without iterative optimization, identifying two lead compounds with sub-micromolar potencies (IC_50_ *<* 1*µ*M and *K*_d_ *<* 500nM) and favorable selectivity profiles with a substantial (*>*50-fold) therapeutic window that decouples on-target metabolic inhibition from non-specific cellular toxicity. Ultimately, HSX establishes a scalable framework that natively integrates generative MPO with automated synthesis, forging a practical path toward fully autonomous drug discovery.

### Hyper Screening X enables synthesis-aware generative MPO for drug design

The HSX framework achieves this synthesis-aware MPO at scale by mapping the reaction logic of the Synple’s automated synthesis platform directly onto the generative agent’s action space (Fig. 1). Collectively, the Synple and eMolecules virtual libraries define a make-on-demand space spanning tens of trillions of compounds, accessed via approximately 90 multi-step synthesis workflows. While these workflow-driven approaches ensure high synthetic feasibility, they introduce strict sequential dependencies such as protection and deprotection steps that conventional generative models relying on flat reaction templates frequently fail to capture. HSX navigates this complexity by formulating the workflow-guided sequential assembly of building blocks as an adaptive Markov decision process (Fig. 1B). Within this structured space, a dynamic transition policy actively penalizes deviations from drug-like constraints, preventing the combinatorial generation of excessively complex molecules and maximizing sampling efficiency. By sampling assembly trajectories proportional to their multi-parameter rewards, the agent enables broad, target-aware exploration to uncover rare, highly diverse hits. Consequently, every generated molecule natively encodes a machine-validated protocol, guaranteeing immediate hardware executability without manual retrosynthetic planning.

**Figure 1:**
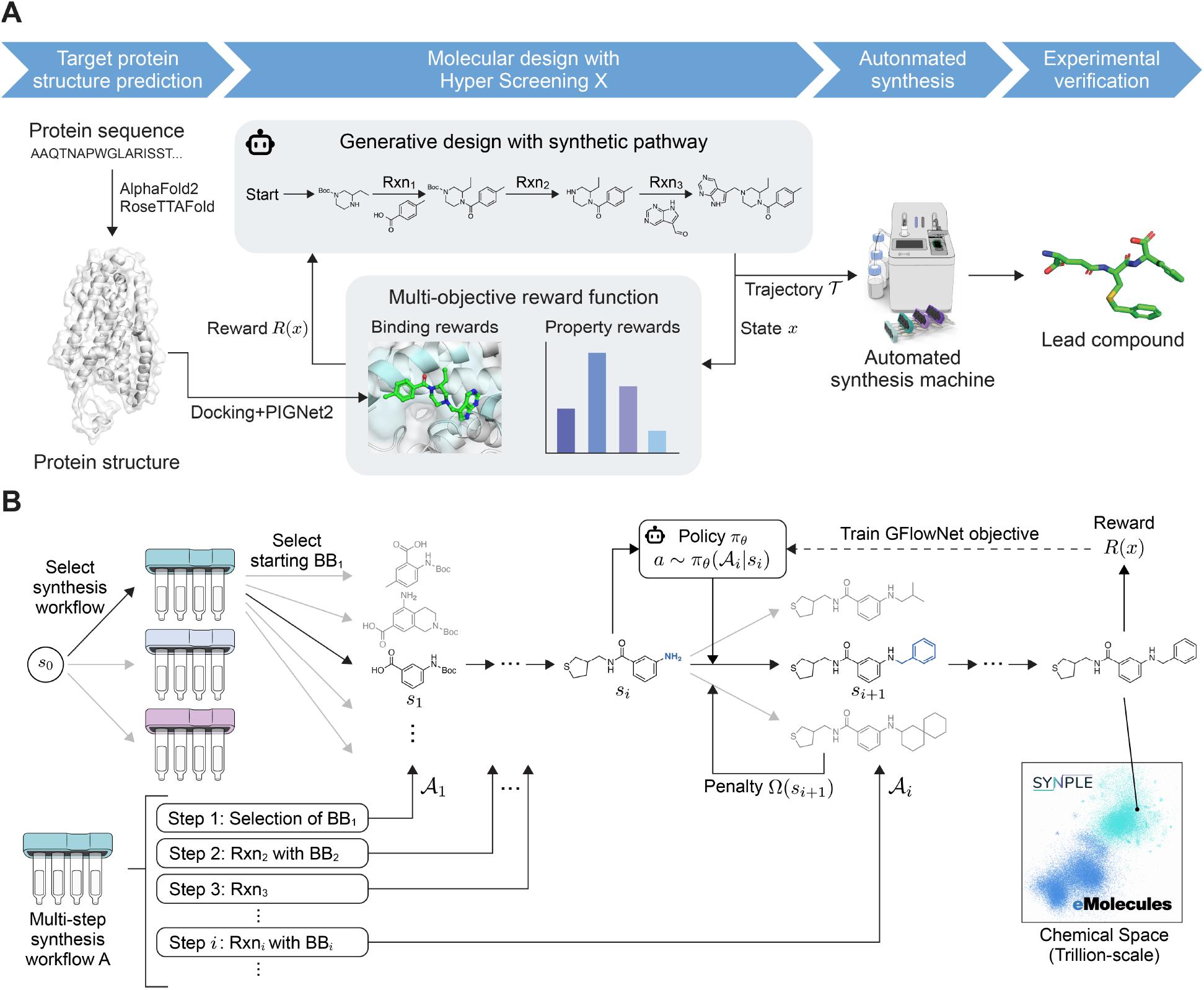
Overview of Hyper Screening X. **A**, Integrated design–make workflow for automated drug discovery. The agent iteratively samples molecules *x* and machine-executable synthetic pathways *T*, evaluates binding affinities and physicochemical properties to compute multi-objective rewards *R*(*x*), and refines the design space. After training, the generated pathways are executed on Synple’s automated synthesis platform. **B**, RxnFlow architecture for Hyper Screening X. The agent constructs molecules through discrete synthetic steps (Rxn_1_, …, Rxn_*n*_), thereby defining an explicit pathway for each molecule. It first selects a multi-step workflow with a predefined action space *A*_*i*_ at each step, ensuring that the product belongs to the Synple and eMolecules make-on-demand libraries and that its pathway is machine-compatible. Actions comprise unimolecular transformations or bimolecular couplings with reactant building blocks. Stochastic transitions follow a learnable policy *π*_*θ*_(*A*_*i*_ |*s*_*i*_) adjusted by the penalty function Ω to promote drug-like space. The terminal molecule is scored by *R*, and the policy is optimized using a generative flow network objective.

During training, the agent simultaneously constructs molecules and their corresponding synthetic pathways, guided by a composite reward integrating PIGNet2-predicted binding affinities (37) and physicochemical constraints (Fig. 1A). The fully trained policy then acts as an autonomous generator of highly synthesizable, property-optimized candidates. Furthermore, the framework is highly modular, meaning that building block libraries and transition parameters can be adjusted *post hoc* to prioritize secondary objectives such as cost, diversity, or specific physicochemical profile without requiring model retraining.

### Generative MPO outperforms virtual screening by orders of magnitude

To benchmark synthesis-aware generative MPO against conventional VS, we conducted an *in silico* campaign targeting microtubule affinity-regulating kinase 4 (MARK4; PDB ID: 5ES1). We established rigorous MPO criteria combining docking score thresholds with four stringent physicochemical constraints: molecular weight (MW) ≤ 500, topological polar surface area (TPSA) ≤ 90, hydrogen-bond donors (HBD) ≤ 4, and octanol–water partition coefficient (LogP) between 3 and 5. Based on random sampling, only ∼ 2% of the underlying make-on-demand library satisfied these four constraints natively. This severely restricted baseline perfectly illustrates the optimization void, where molecules simultaneously possessing high potency and favorable drug-like properties are vanishingly rare, necessitating the intelligent exploration of far larger chemical spaces.

To navigate this void, we trained a single HSX agent across the multi-trillion-scale landscape using a composite reward that jointly optimizes docking affinity and physicochemical properties. We then compared HSX against a brute-force VS baseline comprising one million randomly sampled molecules from the same space. Both approaches generated equally sized evaluation pools, with statistical robustness confirmed via bootstrap resampling.

The performance of HSX was quantified using an enrichment factor (EF) defined as the ratio of valid MPO hits generated by HSX to those found in the random baseline at given docking score thresholds (Fig. 2). Strikingly, the log(EF) scaled linearly as docking thresholds became stricter, demonstrating that the advantage of HSX grows exponentially as design criteria become more demanding. For instance, at a highly stringent threshold where the random baseline yielded 100 hits (EF_100_), HSX identified approximately 1,300-fold more valid candidates on average.

**Figure 2:**
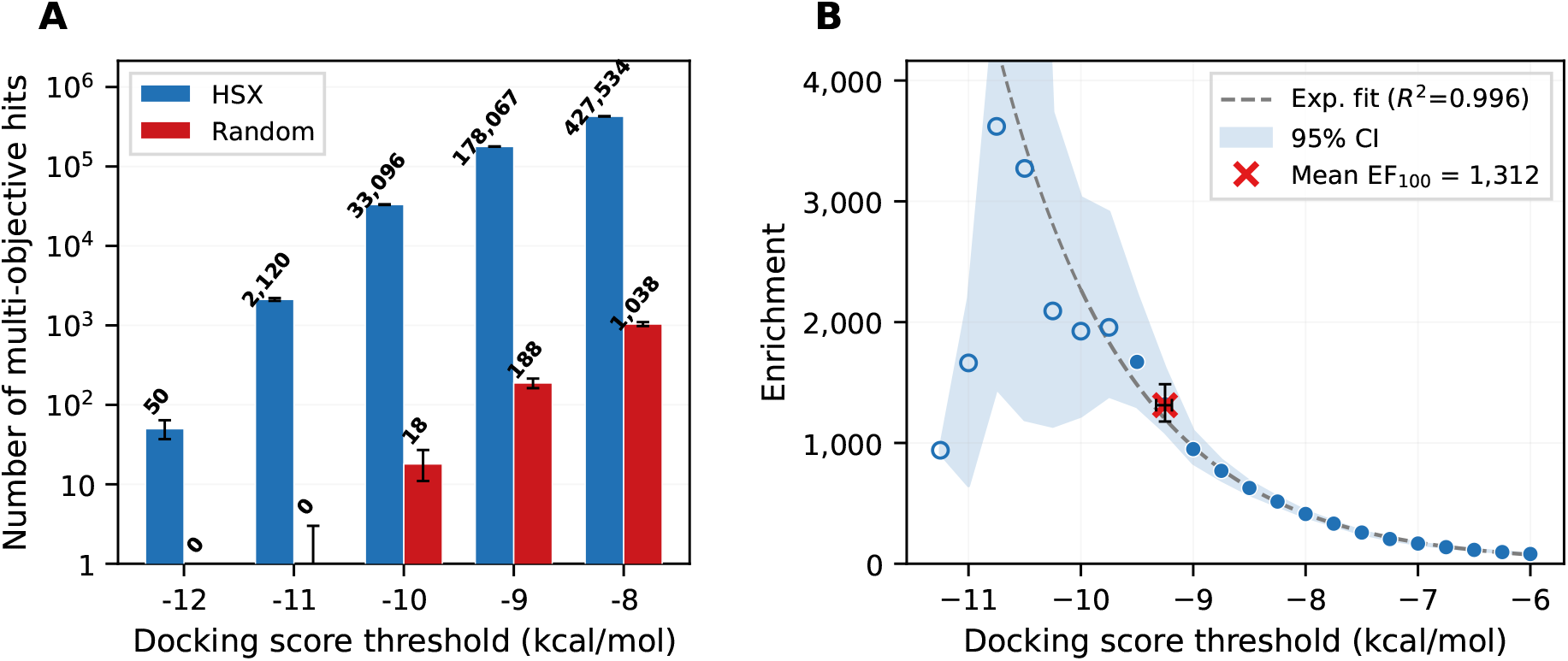
Enrichment of virtual MPO hits identified by HSX. **A**, Number of MPO hits sampled by HSX (blue) and random baseline (red) at each docking score threshold. Both pools were subsampled to one million compounds per bootstrap iteration (total 1,000 iterations). The error bars represent the 95% bootstrap confidence interval. **B**, Enrichment factor (EF) as a function of docking score threshold, defined as the ratio of the MPO hits of the HSX pool to those of the random pool at each threshold. Filled circles indicate thresholds where the random pool contains ≥ 30 hits (stable estimates), while open circles indicate fewer random hits. The dashed line shows an exponential fit to the stable region (*R*^2^ = 0.996), and the shaded band represents the 95% bootstrap confidence interval. The red cross marks EF_100_, the enrichment factor at the threshold that yields 100 random hits, indicating that HSX can identify ∼ 1,300-fold more drug-like, high-potent compounds than the random sampling.

Such substantial and consistent gains reflect HSX’s ability to perform true MPO by jointly encoding all constraints within a unified reward function. By overcoming the sparse nature of MPO hits typical of random enumeration, this goal-aware generative approach substantially outperforms conventional VS, providing a compelling theoretical foundation for the exceptional experimental hit rates observed in subsequent validations.

### Hyper Screening X discovers first-in-class SLC1A5_var inhibitors

Following the *in silico* benchmarking on MARK4, which demonstrated HSX’s capacity to overcome the multi-parameter optimization void, we next sought to evaluate the framework’s prospective capabilities in a more challenging biological context. To this end, we applied it to the SLC1A5 variant (SLC1A5_var), a metabolically critical yet previously undrugged mitochondrial glutamine transporter, as a target protein. We defined a multi-parameter reward function tailored to this target, integrating binding affinity for a cryptic interprotomer site (predicted by PIGNet2 (37)) with key physicochemical constraints (MW ≤ 500, LogP ≤ 5, HBD ≤ 5, HBA ≤ 10). Following the evaluation of 2.2 million molecules during training, we generated approximately 14 million candidates. These candidates were ranked by their reward scores, and the top 23 molecules were selected, excluding those with building block procurement issues such as high pricing or stock unavailability. Executing the AI-generated synthetic routes directly on Synple’s automated synthesis platform yielded 22 successful syntheses, a 95.7% success rate (Fig. 3A). The synthesized molecules were denoted **AI-iMQT-001** through **AI-iMQT-022**, numbered sequentially in order of their rewards.

**Figure 3:**
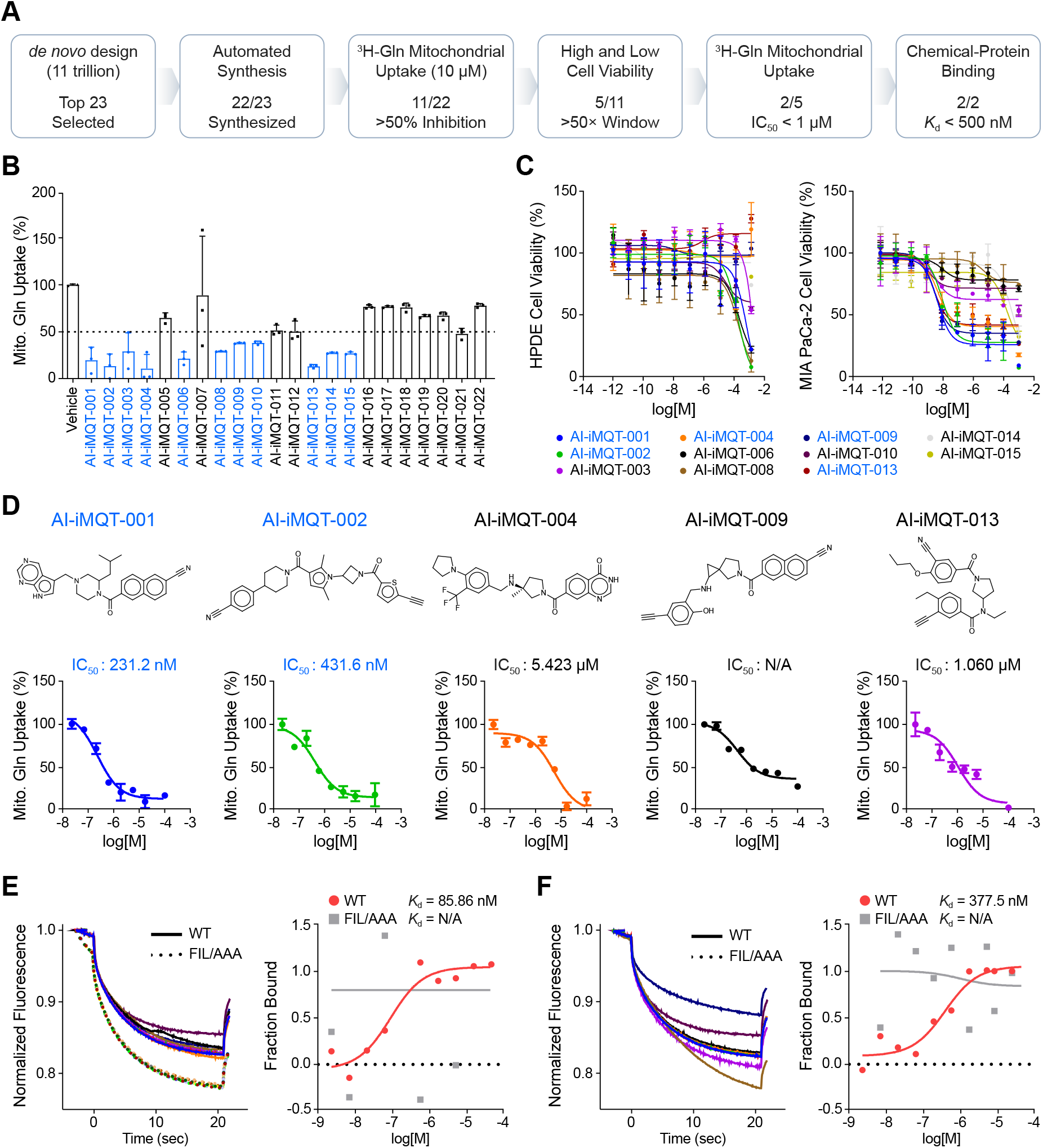
Lead compound identification by HSX. **A**, Screening scheme and selection criteria for lead compounds against SLC1A5_var. **B**, Mitochondrial tritium-labeled glutamine uptake by the 22 HSX-designed molecules synthesized on the Synple platform at 10 *µ*M (n=3 biological replicates). Eleven compounds (blue) showed *>* 50% inhibition. **C**, Cell-viability dose–response curves for SLC1A5_var-high MIA PaCa-2 and SLC1A5_var-low HPDE cells for the 11 primary-screening hits (n=3 biological replicates). The therapeutic window was calculated as the MIA PaCa-2-to-HPDE IC_50_ ratio; five compounds (blue) exceeded 50. **D**, Mitochondrial glutamine-uptake dose–response curves and IC_50_ values for **AI-iMQT-001, -002, -004, -009, and -013** (n=3 biological replicates); calculated values are shown above the plots. **E** and **F**, Microscale thermophoresis (left) and bound fraction (right) for **AI-iMQT-001** (**E**) and **AI-iMQT-002** (**F**) with SLC1A5_var WT (solid line) or FIL/AAA (dotted line). The respective binding constants were 85.86 and 377.5 nM. Data in **B, C, E**, and **F** are mean ± standard deviation.

Functional screening showed that 11 of these 22 compounds inhibited mitochondrial glutamine uptake by *>*50.0% at 10 *µ*M, achieving a 50.0% hit rate without iterative optimization (Fig. 3B). To assess cellular selectivity, we compared IC_50_ values in SLC1A5_var-high (MIA PaCa-2) versus SLC1A5_var-low (HPDE) cell lines (Fig. 3C). Five compounds (**AI-iMQT-001, -002, -004, -009, and -013**) demonstrated a *>*50-fold selectivity window (Table S3). Notably, these hits exhibited substantial structural diversity, spanning five distinct chemotypes, which is a direct consequence of the GFlowNet objective to sample diverse high-reward solutions rather than converging on a single chemotype. The predicted binding poses of the top 2 compounds (**AI-iMQT-001 and -002**) are visualized in Fig. S1.

Further characterization in mitochondrial glutamine uptake assays revealed divergent potency profiles (Fig. 3D). While several hits showed micromolar activity comparable to known inhibitors (38), the two top-ranked compounds, **AI-iMQT-001** and **AI-iMQT-002** achieved sub-micromolar potencies (IC_50_ = 231.2 nM and 431.6 nM, respectively). Their direct target engagements were confirmed by microscale thermophoresis with purified eGFP-tagged SLC1A5_var. Both compounds bound the wild-type protein dose-dependently (*K*_d_ = 85.86 nM and 377.5 nM, respectively) but showed no detectable binding to a binding-deficient FIL/AAA mutant (Figs. 3E, F). These results demonstrate that HSX enables an end-to-end automated workflow from design and synthesis to functional validation, successfully identifying diverse, potent, and target-specific inhibitors for a previously intractable target.

### AI-iMQT-001 and AI-iMQT-002 selectively inhibit SLC1A5_var and disrupt glutamine metabolism

To determine the underlying mechanism of action, we investigated the effects of the two lead compounds on the SLC1A5_var homotrimer. We hypothesized that targeting the cryptic interprotomer interface destabilizes this active complex (38, 39). Size-exclusion chromatography of eGFP-tagged SLC1A5_var showed a dominant trimer peak in vehicle-treated samples. In contrast, treatment with either compound shifted elution profiles toward lower-order species, indicating partial trimer disassembly (Fig. 4A). Co-immunoprecipitation confirmed a substantial reduction in interprotomer association (Fig. 4B). Thus, both **AI-iMQT-001** and **AI-iMQT-002** directly disrupt the trimeric assembly of SLC1A5_var.

**Figure 4:**
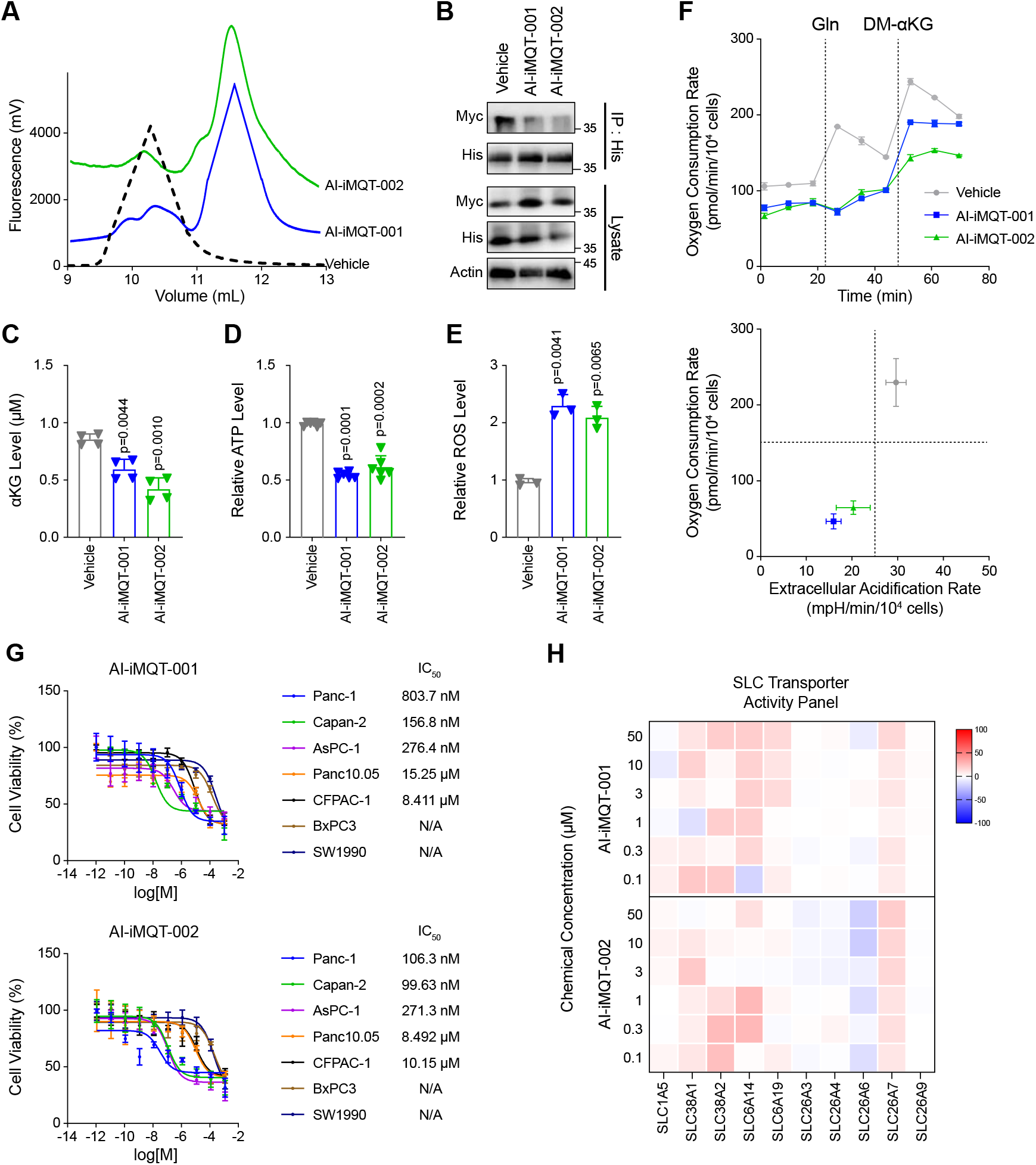
AI-iMQT-001 and AI-iMQT-002 disrupt SLC1A5_var and glutamine metabolism. **A**, Size-exclusion chromatography of eGFP-tagged SLC1A5_var WT after vehicle or 1 *µ*M **AI-iMQT-001 or -002. B**, Co-immunoprecipitation of His- and Myc-tagged SLC1A5_var with His antibody after the same treatments. **C**–**E**, Cellular alpha-ketoglutarate (**C**; n=4), relative ATP (**D**; n=6), and reactive oxygen species (**E**; n=3) after 24 h of vehicle or 1 *µ*M **AI-iMQT-001 or -002**; all n values are biological replicates. **F**, Metabolic phenotype of MIA PaCa-2 cells after the same 24-h treatments (n=3 biological replicates). Top, oxygen consumption after sequential injection of 2 mM glutamine and 2 mM dimethyl-alpha-ketoglutarate; bottom, treatment-dependent metabolic phenotype. **G**, Cell-viability dose–response curves for SLC1A5_var-high Panc-1, Capan-2, and AsPC-1 cells and SLC1A5_var-low Panc10.05, CFPAC-1, BxPC3, and SW1990 cells. **H**, Activity of SLC1A5, SLC38A1, SLC38A2, SLC6A14, SLC6A19, SLC26A3, SLC26A4, SLC26A6, SLC26A7, and SLC26A9 across concentrations of **AI-iMQT-001 and -002**. Activities are normalized to control (0%) and vehicle (100%; n=3). Data in **C**–**G** are mean ± standard deviation.

Functionally, this structural disruption depleted intracellular *α*-ketoglutarate (*α*KG) and ATP while increasing reactive oxygen species, hallmarks of impaired glutamine-dependent anaplerosis (39) (Figs. 4C–E). Seahorse Metabolic flux analysis revealed a pronounced reduction in a basal oxygen consumption rate (OCR). Exogenous glutamine failed to rescue mitochondrial respiration, whereas supplementation with dimethyl-*α*-ketoglutarate (DM-*α*KG) restored OCR (Fig. 4F). This indicates a specific glutaminolysis blockade, establishing that SLC1A5_var trimer disruption precipitates mitochondrial metabolic collapse.

Compound efficacy tightly correlated with target abundance across a panel of pancreatic cancer cell lines. Both compounds exhibited nanomolar IC_50_ values in SLC1A5_var-high-expressing lines (Panc-1, Capan-2, AsPC-1), with drastically reduced potency and negligible toxicity in low-expressing lines (Panc10.05, CFPAC-1, BxPC3, SW1990) (Fig. 4G). Profiling against related SLC transporters and ion channels confirmed minimal off-target activity (Fig. 4H). Collectively, these compounds selectively engage SLC1A5_var, disrupt its functional trimeric state, and induce targeted metabolic collapse, defining an unprecedented mechanism for inhibiting mitochondrial glutamine transport.

## Discussion

The realization of fully autonomous drug discovery has long been stymied by the fundamental disconnect between expansive virtual design and physical executability. HSX provides a definitive solution to this challenge, representing a fundamental paradigm shift from the retrospective filtering of static, billion-scale libraries to the active, prospective generation of synthesis-guaranteed molecular matter.

Crucially, our empirical results explicitly validate the concept of the optimization void. We demonstrate that navigating a multi-trillion-scale landscape is not merely advantageous, but statistically imperative to discover rare molecules that simultaneously satisfy stringent MPO profiles. By directly embedding the deterministic reaction logic of automated synthesis into the GFlowNet architecture, HSX circumvents the combinatorial explosion of chemically intractable structures. This continuous integration of algorithmic exploration and hardware-compatible execution translates into an exceptional real-world performance: an unprecedented 96% experimental synthesis success rate (22 out of 23), a 50% functional hit rate, far eclipsing the yield of conventional generative or virtual screening methodologies.

The true measure of any *de novo* design framework, however, lies in its capacity to drug challenging biological targets. The prospective identification of two lead compounds, **AI-iMQT-001** and **AI-iMQT-002**, underscores the practical supremacy of this approach. By successfully targeting a cryptic interprotomer interface in SLC1A5_var, a topological mechanism notoriously elusive to conventional screening, HSX proved its ability to navigate complex structural demands. The resulting leads not only exhibit sub-micromolar potency but also maintain a broad therapeutic window, cleanly decoupling on-target metabolic inhibition from nonspecific cytotoxicity.

Taken together, these findings firmly establish synthesis-aware generative AI as a transformative engine for early-stage drug discovery. By resolving the algorithmic and synthetic bottlenecks that currently limit chemical exploration, HSX directly bridges the gap between *in silico* design and experimental validation. Ultimately, this framework lays a robust and actionable foundation for the fully autonomous, closed-loop discovery ecosystems of the future.

## Acknowledgments

This work was supported by National Research Foundation of Korea (NRF) funded by the Korean government (MSIT) (NRF-2023R1A2C2004376, RS-2023-00257479 to W.Y.K; RS-2025-00573098, RS-2025-18362970, and RS-2026-25510253 to J.M.H.). It was also supported by the InnoCORE program of the Ministry of Science and ICT (N10260004 to W.Y.K) and Yuhan Innovation Project (to J.M.H.).

## Supplementary Materials

### A Hyper Screening X architecture

#### A.1 Synthesis space and generative formulation

The molecular generation engine of HSX is based on the RxnFlow model (22), reconfigured to operate on Synple’s automated synthesis platform using Synple and eMolecules virtual libraries (36). The two libraries are defined by approximately 90 reaction workflows composed of commercially available building blocks, spanning a chemical space of tens of trillions of virtual compounds. The libraries are continually expanded as new building blocks and reactions are integrated, leading to a steadily growing chemical space. Each reaction workflow prescribes a fixed sequence of two or three synthetic protocols consisting of an initial building-block selection, optional unimolecular transformations (e.g., ring closures or protecting-group manipulations), and bimolecular coupling reactions, together with the permissible building-block categories for each step. Because every generative trajectory strictly follows one of these predefined protocols, all output molecules are by construction executable on the Synple platform without additional retrosynthesis planning. HSX can be accessed on the HyperLab platform (40). The source code is available at https://github.com/SeonghwanSeo/RxnFlow.

HSX formulates molecule generation as a hierarchical Markov decision process in which the agent first selects a reaction workflow and then sequentially selects building blocks from the categories dictated by the selected workflow. The reward is calculated only for the molecules in the terminating state. The forward policy is trained according to the trajectory balance objective (41) with a uniform backward policy. This greatly simplifies the learning problem. To promote structural diversity, a random workflow strategy uniformly samples reaction workflows during training.

Two mechanisms control the effective action space at each building-block selection step. First, because the building-block library is highly imbalanced across categories, we apply an action-space subsampling strategy (22). Building blocks are stratified into five clusters based on commercial availability, and an arbitrary fraction is sampled uniformly from each cluster at every training step, with importance weights log(1*/*sampling ratio) correcting the resulting bias in transition probability. Second, we introduced a penalty function, Ω(·), to constrain compositional physicochemical properties that accumulate incrementally during molecular assembly, such as molecular weight and total atom count. If the projected molecular size of a subsequent state exceeds a defined constraint threshold, the corresponding action is penalized within the forward transition probability.

#### A.2 Model architecture

The HSX generative policy is parameterized by a graph neural network that maps the current molecular intermediate to a distribution over available actions. The policy network is trained using trajectory balance (TB) objective (41), which is a standard GFlowNet training objective for scientific discovery tasks. To prevent mode collapse, we adopted the double GFlowNet (DGFN) training strategy (42) by using an exponential moving average with a decay rate of 0.99.

##### State encoder

To capture the structural features of the evolving intermediate, we encode its molecular graph using a directed graph transformer architecture (43). Node attributes include one-hot encodings of atom type, chirality, formal charge, aromaticity, explicit hydrogen count, and isotope labels; edge features encode the formal bond types. To guide the generative process through the multi-objective optimization landscape, global attributes include 9 physicochemical properties of the state molecule: heavy-atom count, hydrogen bond acceptors (HBA), hydrogen bond donors (HBD), ring count, aromatic ring count, rotatable bonds, molecular weight (MW), topological polar surface area (TPSA), and octanol-water partition coefficient (log *P*). We use a global graph representation with the final graph-level readout.

##### Reaction encoder

Each reaction type is represented with a learnable embedding vector. We note that the same reactions are distinguished depending on which multi-step synthesis workflow it belongs to.

##### Reactant encoder

Each candidate building block of each reaction is represented by four components: (i) a molecular fingerprint comprising Morgan fingerprints (44) (radius 2, 1024 bits) concatenated with MACCS keys (166 bits) (45); (ii) normalized physicochemical properties (the same 9 descriptors as above); (iii) a learnable reactant-type embedding; and (iv) a learnable synthetic-accessibility tier embedding (5 tiers). These four representations are concatenated and projected to yield a dense reactant embedding.

##### Policy heads

The policy network maps the current state embedding to transition probabilities through a sequentially structured dual-head framework: a synthetic workflow selection head followed by a reactant selection head. The synthetic workflow selection head first maps the global graph embedding to produce a probability distribution over the pre-defined synthetic pathways (workflow). The reactant selection head subsequently conditions the global graph embedding on a learnable embedding representing the chosen reaction template via element-wise addition, projecting the result through an MLP to match the 64-dimensional reactant embedding space. The reactant selection transition probabilities are then computed as the inner product of this conditioned state representation with the candidate reactant embeddings. To calibrate exploration across varying target stringencies, we scale these raw logits using a learnable temperature-conditional parameter that dynamically modulates the sharpness of the policy distribution with respect to the continuous temperature condition *β* (46).

##### Partition function head

To train the model using a trajectory-balance objective (41), the model also predicts a log-partition-function log *Z*(*β*) for a given temperature condition *β*.

#### A.3 Action-space subsampling

At each reactant selection step, the full building-block library for the relevant category may contain thousands to tens of thousands of candidates, making exhaustive scoring computationally expensive. To manage this, we introduces the action subsampling strategy (22). HSX stratifies building blocks within each category into five clusters defined by their commercial availability (price tier from the eMolecules catalog). At every training step, a fixed fraction of blocks is sampled uniformly from each cluster, subject to a minimum of 50 blocks per cluster. The resulting non-uniform inclusion probabilities are corrected during policy evaluation by adding importance weights *w* = log(1*/r*_*c*_) to the log-probabilities, where *r*_*c*_ is the sampling ratio for cluster *c*. This action subsampling strategy introduces bias into the partition function estimation, but enables unbiased estimation for the forward transition probability during inference.

#### A.4 Reward function

The composite reward for each generated molecule is defined as the product of a docking-affinity reward and a physicochemical-property penalty;

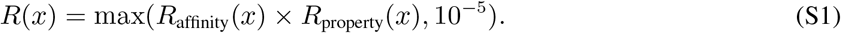

where *x* is the termination state of the generative trajectory. The reward is clipped to 10^−5^ for GFlowNet training, which requires positive rewards.

##### Affinity reward

Binding poses are generated by GPU-accelerated UniDock (47) and scored by the PIGNet2 model (37).

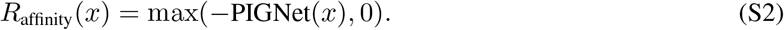

##### Property reward

For each physicochemical property *p* with an allowed range [*p*_min_, *p*_max_] and a smoothness margin *σ*, the penalty is

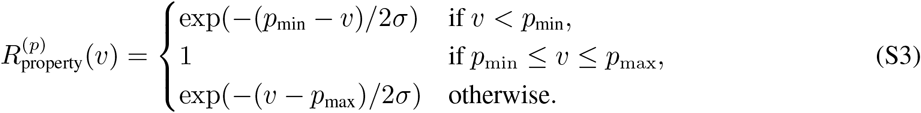

The overall property reward is the product over all constrained properties: 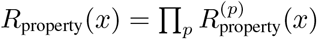 This smooth penalty preserves the training gradient near constraint boundaries rather than imposing hard binary cutoffs that silence the learning signal. The margin parameter *σ* was set to 50.0 for MW, 1.0 for LogP, 1 for HBD, and 2 for HBA.

#### A.5 Property-level penalty function

For compositional physicochemical properties whose molecular values are approximately the sum of their building-block contributions (e.g., MW, HBD, HBA, TPSA), HSX enforces property constraints directly within the generative process. Each constrained property *p* is assigned a threshold *θ*_*p*_ (e.g., *θ*_MW_ = 500; *θ*_HBD_ = 5). At each decision step, we compute the penalty function Ω(*s*_*i*+1_) based on the next intermediate state *s*_*i*+1_ as:

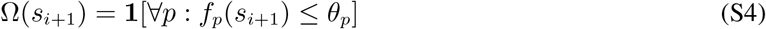

This mechanism structurally prevents the generation of property-violating molecules without requiring post-hoc filtering.

### B Molecular generation pipeline

#### B.1 Preparation of the structure of SLC1A5_var

The structure of SLC1A5_var was prepared as previously described (38). Briefly, the monomeric structure was constructed using RoseTTAFold2.0 via ColabFold on a local GPU cluster using the query sequence without templates or multiple sequence alignments (48, 49). The resulting monomer was assembled into a homotrimer using the MZ-Dock server (50), targeting the transmembrane domain 3 as the interprotomer interaction site, and it was subsequently refined through energy minimization and structural relaxation via Desmond molecular dynamics simulation (51). This refined trimer was then processed using the protein preparation wizard in Maestro (version 9.6, Schrödinger, LLC) to assign bond orders, add missing hydrogens, and optimize a hydrogen-bond network, followed by restrained minimization using the OPLS3 force field with a maximum root mean square distance of 0.30 Å(52). The receptor grid box (20 Å × 20 Å × 20 Å) centered on the centroid of residues Phe97, Ile104, and Leu105, which is the site of the interprotomer interaction of SLC1A5_var, was generated to define the putative binding pocket.

#### B.2 Training of Hyper Screening X

To design inhibitors targeting the MQT homotrimer, we trained the HSX model using a multi-parameter reward function that simultaneously optimizes target binding affinity and physicochemical properties. During training, we employed GPU-accelerated UniDock (47) to compute binding pose candidates of the generated molecules. We evaluated the poses using the PIGNet2 model (37), where the predicted binding affinities serve as one of the rewards together with physicochemical properties for training. The center of the search box for docking was set to the center coordinate of the binding site, with a box size of 20 Å × 20 Å × 20 Å. In addition, we incorporated four physicochemical property criteria into the multi-parameter reward function according to Lipinski’s rule of five: MW ≤ 500, LogP ≤ 5, HBD ≤ 5, and HBA ≤ 10. Rather than employing binary filters that yield zero reward for TPP violations and effectively silence the training signal, we implemented a smooth penalty reward function with permissive margins. This approach down-weights, but does not strictly forbid, trajectories that slightly violate the desired window, thereby facilitating optimization. Lastly, to maximize the structural diversity of the generated molecules, we trained models in parallel, both with and without the random workflow strategy, and utilized all resulting models for molecule generation.

**Table S1:** Training hyperparameters for the HSX model.

| Parameter | Value |
| --- | --- |
| Online samples per iteration | 64 |
| Replay samples per iteration | 128 |
| Random exploration probability ($\varepsilon$) | 0.1 |
| Learning rate | $10^{-4}$ |
| Learning rate ($\log Z_\theta$) | $10^{-2}$ |
| Reward exponent ($\beta$) | Uniform [1.0, 64.0] |
| Action subsampling ratio | 0.1 |
| Graph transformer layers | 4 |
| Graph transformer dimension | 256 |
| Graph transformer heads | 4 |
| State embedding dimension | 512 |
| Block embedding dimension | 64 |
| Dropout | 0.1 |

#### B.3 Molecule generation by Hyper Screening X and candidate selection

We generated molecules from the trained models and removed duplicates, yielding ∼ 14 million unique molecules. These molecules were strictly filtered to satisfy all physicochemical-property conditions included in the multiparameter training. To prevent bias toward specific building blocks within this pool, we applied a diversity control algorithm that restricted the maximum usage frequency of any individual building block to 100, which narrowed the pool down to ∼ 1.4 million molecules. After filtering the molecules at the 2D-structure level as above, we employed Hyper Binding from HyperLab to accurately predict binding affinities (40). Hyper Binding combines an exhaustive search method for 3D conformational exploration with the PIGNet2 model (37) for binding-affinity prediction. Based on the Hyper Binding results, we selected molecules whose docked conformers (i) formed seven protein–ligand interactions considered important for target activity (hydrogen bonding with R263; *π*–*π* interaction with F97; hydrophobic interactions with V83, I87, M90, V93 and L260), and (ii) had a predicted binding affinity of ≤ -6 kcal/mol. Next, for cost-efficiency, we restricted the price of the building blocks, further reducing the set to 130 molecules. We then applied medicinal chemistry filters to exclude structures expected to exhibit poor drug-likeness, specifically by limiting the number of acetylenes and methyl alkynes, yielding 105 molecules. From this final pool, we ultimately selected 23 molecules prioritized by the predicted binding affinities while also considering building-block costs.

**Figure S1:**
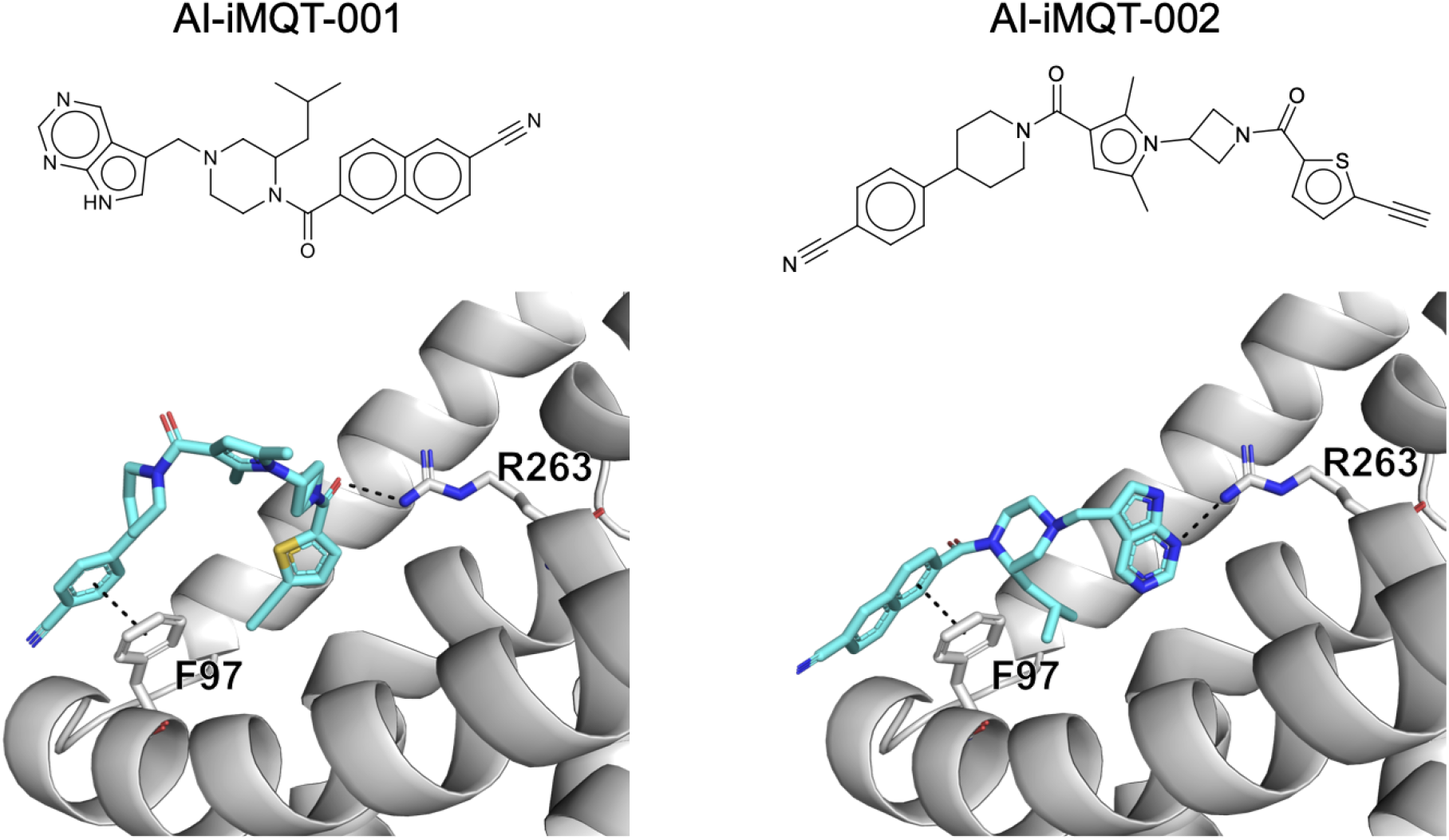
Predicted binding pose of AI-iMQT-001 and AI-iMQT-002. Both molecules exhibit *π*–*π* stacking interactions with F97 and form hydrogen bonds with R263.

**Table S2:**
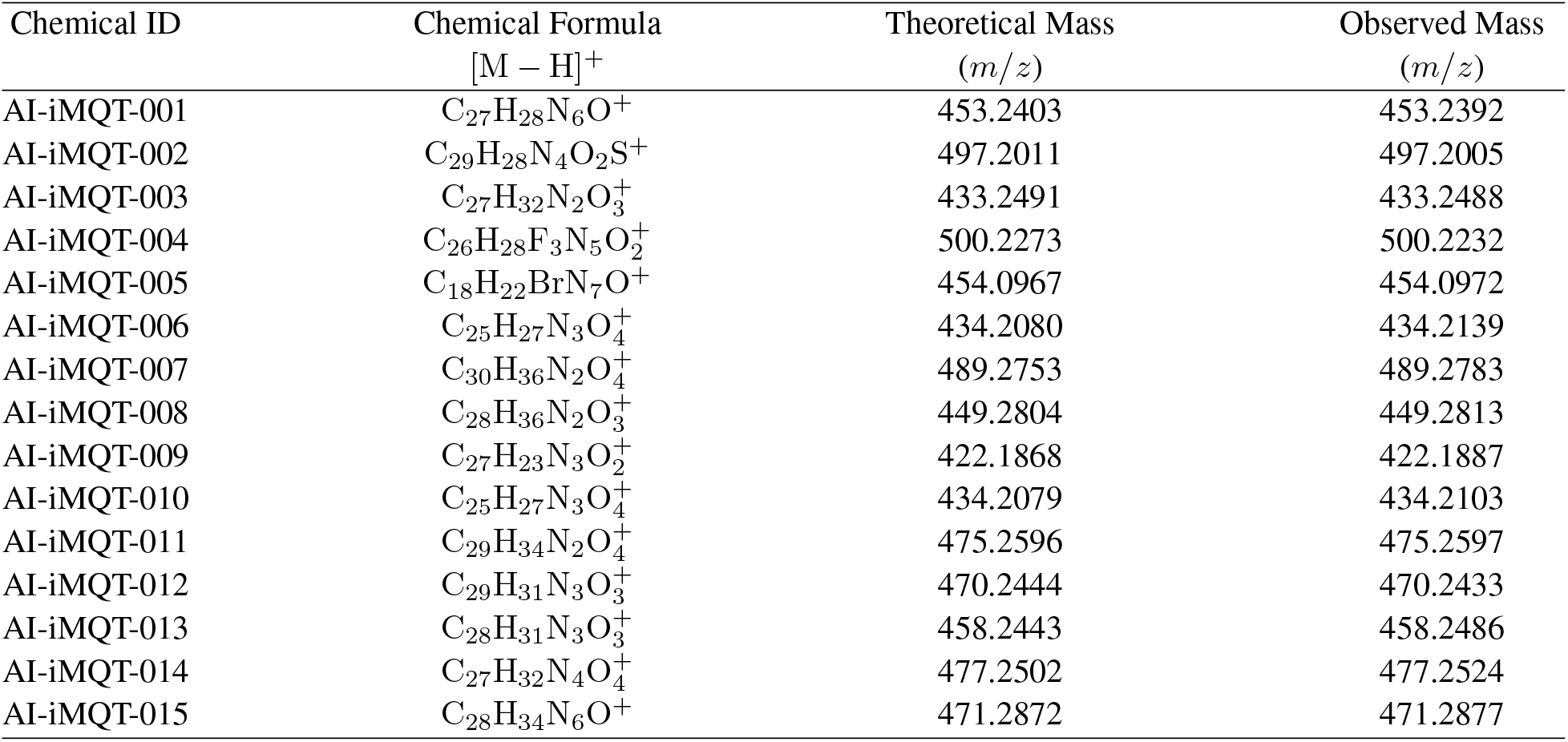
High resolution mass spectrometry of compounds tested. Calculated and exact found mass detected through HRMS.

| Chemical ID | Chemical Formula<br>[M - H]<sup>+</sup> | Theoretical Mass<br>(*m/z*) | Observed Mass<br>(*m/z*) |
| --- | --- | --- | --- |
| AI-iMQT-001 | C<sub>27</sub>H<sub>28</sub>N<sub>6</sub>O<sup>+</sup> | 453.2403 | 453.2392 |
| AI-iMQT-002 | C<sub>29</sub>H<sub>28</sub>N<sub>4</sub>O<sub>2</sub>S<sup>+</sup> | 497.2011 | 497.2005 |
| AI-iMQT-003 | C<sub>27</sub>H<sub>32</sub>N<sub>2</sub>O<sub>3</sub><sup>+</sup> | 433.2491 | 433.2488 |
| AI-iMQT-004 | C<sub>26</sub>H<sub>28</sub>F<sub>3</sub>N<sub>5</sub>O<sub>2</sub><sup>+</sup> | 500.2273 | 500.2232 |
| AI-iMQT-005 | C<sub>18</sub>H<sub>22</sub>BrN<sub>7</sub>O<sup>+</sup> | 454.0967 | 454.0972 |
| AI-iMQT-006 | C<sub>25</sub>H<sub>27</sub>N<sub>3</sub>O<sub>4</sub><sup>+</sup> | 434.2080 | 434.2139 |
| AI-iMQT-007 | C<sub>30</sub>H<sub>36</sub>N<sub>2</sub>O<sub>4</sub><sup>+</sup> | 489.2753 | 489.2783 |
| AI-iMQT-008 | C<sub>28</sub>H<sub>36</sub>N<sub>2</sub>O<sub>3</sub><sup>+</sup> | 449.2804 | 449.2813 |
| AI-iMQT-009 | C<sub>27</sub>H<sub>23</sub>N<sub>3</sub>O<sub>2</sub><sup>+</sup> | 422.1868 | 422.1887 |
| AI-iMQT-010 | C<sub>25</sub>H<sub>27</sub>N<sub>3</sub>O<sub>4</sub><sup>+</sup> | 434.2079 | 434.2103 |
| AI-iMQT-011 | C<sub>29</sub>H<sub>34</sub>N<sub>2</sub>O<sub>4</sub><sup>+</sup> | 475.2596 | 475.2597 |
| AI-iMQT-012 | C<sub>29</sub>H<sub>31</sub>N<sub>3</sub>O<sub>3</sub><sup>+</sup> | 470.2444 | 470.2433 |
| AI-iMQT-013 | C<sub>28</sub>H<sub>31</sub>N<sub>3</sub>O<sub>3</sub><sup>+</sup> | 458.2443 | 458.2486 |
| AI-iMQT-014 | C<sub>27</sub>H<sub>32</sub>N<sub>4</sub>O<sub>4</sub><sup>+</sup> | 477.2502 | 477.2524 |
| AI-iMQT-015 | C<sub>28</sub>H<sub>34</sub>N<sub>6</sub>O<sup>+</sup> | 471.2872 | 471.2877 |

## C Experimental validation

### C.1 Synthesis and characterization of compounds

High-resolution mass spectrometry (HRMS) data is provided in Table S2.

### C.2 Purification of mitochondria from cells and mitochondrial amino acid uptake assay

The mitochondria were prepared according to the methods of Wieckowski et al., with slight modification (53). For each experiment, MIA PaCa-2 cells that were stably transfected with 3XMyc-EGFP-OMP25 were immunoprecipitated with an anti-Myc antibody. Then, 100 *µ*l of anti-Myc magnetic beads (Pierce) was prewashed three times with KPBS. All washes were performed by gentle pipetting via a wide-bore pipette tip, and the beads were collected with a magnetic stand (DynaMag Spin Magnet, Invitrogen). A total of 1 × 10^7^ cells cultured in 15-cm dishes were washed twice with DPBS and then gently scraped into 1 mL of KPBS. The cells were subsequently centrifuged at 900 × *g* for 3 min, after which the pellet was resuspended in 1 mL of KPBS. The cell mixture was carefully homogenized with 50 strokes in a 2-mL Dounce homogenizer to avoid air bubbles. The homogenate was centrifuged at 1, 000 × *g* for 2 min, and the supernatant was immunoprecipitated with anti-c-Myc antibody-conjugated magnetic beads for 20 min. The c-Myc-tagged mitochondria were eluted with 250 mL of 0.25 mg/mL c-Myc peptide (Pierce). Mitochondrial amino acid uptake was measured by first diluting the purified mitochondria in 50 *µ*L of KPBS. This solution was placed in a 1.5-mL tube with vehicle or chemicals in respective concentrations for 24 hours. Radiolabeled glutamine (L-[^3^H]Gln) was used to assess mitochondrial amino acid uptake, followed by incubation at 37 °C for one hour. Uptake was stopped by incubating the tubes at 4 °C. Then, the tubes were centrifuged at 12, 000 × *g* for 15 min, and the supernatant was removed and washed with KPBS twice. The cells were extracted with lysis buffer (0.2% SDS) and mixed with 3 mL of AquaLight+ (HIDEX). All measurements were analyzed by a liquid scintillation counter (HIDEX). The lysate was subsequently used to measure the mitochondrial protein concentration of each sample with a BCA assay kit (Intron). Amino acid uptake was calculated from the counts per minute per sample and the specific activity of each labeled amino acid, and the uptake of these amino acids was normalized to the mitochondrial protein content.

**Table S3:** IC50 of cell viability of Top 11 chemicals on SLC1A5_var low and high expressing pancreatic cancer cell lines. Cell viability IC50 of HPDE and MIA PaCa-2 of 11 chemicals with high mitochondrial glutamine uptake inhibition rate.

| Chemical ID | HPDE (SLC1A5_var Low)<br>IC<sub>50</sub> [μM] | MIA PaCa-2 (SLC1A5_var High)<br>IC<sub>50</sub> [nM] |
| --- | --- | --- |
| AI-iMQT-001 | 617.3 | 530.8 |
| AI-iMQT-002 | 131.9 | 108.0 |
| AI-iMQT-003 | NA | NA |
| AI-iMQT-004 | NA | 594.0 |
| AI-iMQT-006 | 161.6 | NA |
| AI-iMQT-008 | 141.1 | NA |
| AI-iMQT-009 | NA | 353.8 |
| AI-iMQT-010 | NA | NA |
| AI-iMQT-013 | NA | 298.8 |
| AI-iMQT-014 | 317.5 | 3697.1 |
| AI-iMQT-015 | 74.25 | 1619.5 |

### C.3 Cell viability and growth measurements

For the cell viability assays, cells were seeded into 96-well culture plates at a density of 10,000 cells per well in the appropriate medium supplemented with 10% dialyzed FBS. After successful cell attachment, the medium was replaced with conditioned medium. After 48 hours, cell viability was measured via a WST-1 assay. The absorbance of each sample at 450 nm (reference wavelength 625 nm) was measured with a microplate reader (Infinite 200 PRO, Tecan) and normalized to that of the control.

### C.4 Microscale thermophoresis

Microscale thermophoresis experiments were performed with Monolith NT.115 (NanoTemper Technologies). eGFP-tagged SLC1A5_var proteins were purified as previously (38). The protein samples were diluted with 20 mM HEPES (pH 7.5), 100 mM NaCl, and 0.05% DDM and assayed with Monolith to confirm the absence of aggregation, revealing the optimal level of initial fluorescence. A series of dilutions with the same buffer were performed as the instrument guided, and each concentration of **AI-iMQT-001** or **AI-iMQT-002** was mixed with the protein at a ratio of 1:1 and incubated inside the Premium Capillary (Nanotemper, MO-K025). Red fluorescence was selected as the LED source, and 50% infrared-laser power was used at 25 °C. Analysis of the data was performed with MO Affinity Analysis 2.3.

### C.5 Coimmunoprecipitation and immunoblotting

Myc-tagged and His-tagged SLC1A5_var WT were co-transfected at a ratio of 1:1. After twenty-four hours, the cells were harvested via lysis buffer (40 mM HEPES (pH 7.4), 0.5% Triton X-100, 10 mM *β*-glycerol phosphate, 10 mM pyrophosphate, and 2.5 mM MgCl_2_) supplemented with a protease inhibitor (5 *µ*g/mL aprotinin, 10 *µ*g/mL leupeptin, and 250 *µ*M PMSF). The lysate was subsequently sonicated on ice. The cell lysates were incubated for 4 hours with 1 *µ*g of the appropriate antibodies in the presence of Protein A/G agarose beads (InvivoGen) at 4 °C. The resulting complexes were washed four times with lysis buffer and eluted with 2 × Laemmli buffer with vigorous vortexing. The eluates were isolated through centrifugation and subjected to SDS–PAGE for immunoblotting. A total of 30 *µ*g of protein was subjected to SDS–PAGE and transferred to a PVDF membrane (Merck) via the Trans-BlotR Turbo Blotting System (Bio-Rad). The membranes were subsequently blocked in TBST buffer containing 2% skim milk and incubated with the appropriate primary antibodies for 18 hours. The membranes were incubated with the appropriate secondary antibodies (anti-mouse or anti-rabbit IgG conjugated with horseradish peroxidase, Invitrogen) for 1 hour. The immunoblot signals were detected with a MicroChemi instrument (DNR Bioimaging Systems) with enhanced chemiluminescence (EzWestLumi, ATTO).

### C.6 Measurement of *α*KG, ATP, and ROS

*α*KG levels were determined with an alpha-ketoglutarate assay kit (ab83431, Abcam) according to the manufacturer’s instruction. Cells seeded in a 24-well plate were treated with **AI-iMQT-001** and **AI-iMQT-002**, lysed using a lysis reagent (Promega), and briefly centrifuged at 13, 000 × *g* for 1 min. ATP was measured by mixing each supernatant and CellTiter-Glo (G7570, Promega) in 1:1 ratio. The mixture was incubated in dark for 10 min and the luminescence was quantified by a plate reader (Infinite 200 PRO, TECAN). Intracellular ROS levels were determined by staining cells with 1 mM 5-(and-6)-carboxy-2’,7’-dichlorodihydrofluorescein diacetate (H2DCFDA, Invitrogen) for 10 min according to the manufacturer’s protocol (Ex/Em: 492/517 nm). The stained cells were analyzed via flow cytometry (BD FACSAria III).

### C.7 OCR/ECAR measurement

The OCR and ECAR were determined with an XFe24 extracellular flux analyzer (Agilent Technologies) as described in the manufacturer’s protocol. A total of 4.0 × 10^4^ cells were seeded per well in 24-well micro cell culture plates for Seahorse (Agilent Technologies) in DMEM supplemented with 10% FBS with vehicle, **AI-iMQT-001**, or **AI-iMQT-002** and incubated at 37 °C in a 5% CO_2_ incubator overnight. Then, the DMEM was removed and replaced with bicarbonate-free DMEM containing phenol red (pH 7.4), and the cells were incubated at 37 °C in a non-CO_2_ incubator to equilibrate the CO_2_ concentration in the atmosphere for one hour. Using the XFe24 analyzer, the OCR and ECAR were measured under baseline conditions and upon treatment with glutamine (Gln, 2 mM) and dimethyl-alpha-ketoglutarate (DM-*α*KG, 2 mM). For the metabolic phenotype assay, the basal OCR and basal ECAR of the cells were measured. The OCR and ECAR values were normalized to the number of cells and analyzed with WAVE software (Agilent Technologies).

### C.8 Transporter and channel protein activity test

SLC1A5-, SLC38A1-, SLC38A2-, SLC6A14-, SLC6A19-, SLC26A3-, SLC26A4-, SLC26A6-, SLC26A7-, or SLC26A9- and YFP-F46L/H148Q/I152L-expressing cells were plated in 96-well culture plates at a density of 2 × 10^4^ cells/well and incubated for 24 hours. Each well of the 96-well culture plates was washed 2 times with 100 *µ*L of DPBS, and the wells were filled with 50 *µ*L of HEPES-buffered solution. Each compound (vehicle, **AI-iMQT-001, AI-iMQT-002**) was added at 0.03, 0.01, 0.3, 0.1, 1, 3, 10, and 50 *µ*M before activity measurement. The 96-well culture plates were placed on a FLUOstar Omega Microplate Reader (BMG Labtech, Ortenberg, Germany) for fluorescence assays. Each well was assayed individually for iodide ion influx mediated by the control treatment or SLC1A5, SLC38A1, SLC38A2, SLC6A14, SLC6A19, SLC26A3, SLC26A4, SLC26A6, SLC26A7, and SLC26A9 by continuously recording the fluorescence (1 second per point) for 5 seconds (baseline). Then, 50 *µ*L of NaI- or NaSCN-substituted HEPES-buffered solution (with NaI or NaSCN replacing NaCl from the original HEPES buffer) was added to the liquid injector at 5 seconds, and the YFP fluorescence was recorded for 10 seconds. The activity of each transporter or channel was calculated by the gradient of the YFP fluorescence quenching at the moment of halide salt injection.

